# *In utero* diffusion tensor imaging of the fetal brain: a reproducibility study

**DOI:** 10.1101/132704

**Authors:** András Jakab, Ruth O`Gorman Tuura, Christian Kellenberger, Ianina Scheer

## Abstract

Our purpose was to evaluate the within-subject reproducibility of i*n utero* diffusion tensor imaging (DTI) metrics and the visibility of major white matter structures.

Images for 30 fetuses (20-33. postmenstrual weeks, normal neurodevelopment: 6 cases, cerebral pathology: 24 cases) were acquired on 1.5T or 3.0T MRI. DTI with 15 diffusion-weighting directions was repeated three times for each case, TR/TE: 2200/63 ms, voxel size: 1*1 mm, slice thickness: 3-5 mm, b-factor: 700 s/mm^2^. Reproducibility was evaluated from structure detectability, variability of DTI measures using the coefficient of variation (CV), image correlation and structural similarity across repeated scans for six selected structures. The effect of age, scanner type, presence of pathology was determined using Wilcoxon rank sum test.

White matter structures were detectable in the following percentage of fetuses in at least two of the three repeated scans: corpus callosum genu 76%, splenium 64%, internal capsule, posterior limb 60%, brainstem fibers 40% and temporooccipital association pathways 60%. The mean CV of DTI metrics ranged between 3% and 14.6% and we measured higher reproducibility in fetuses with normal brain development. Head motion was negatively correlated with reproducibility, this effect was partially ameliorated by motion-correction algorithm using image registration. Structures on 3.0 T had higher variability both with- and without motion correction.

Fetal DTI is reproducible for projection and commissural bundles during mid-gestation, however, in 16-30% of the cases, data were corrupted by artifacts, resulting in impaired detection of white matter structures. To achieve robust results for the quantitative analysis of diffusivity and anisotropy values, fetal-specific image processing is recommended and repeated DTI is needed to ensure the detectability of fiber pathways.

**Abbreviations:**

AD: axial diffusivity;
CCA: corpus callosum agenesis;
CV: coefficient of variation,
DTI: diffusion tensor imaging;
FA: fractional anisotropy;
GW: gestational week;
MD: mean diffusivity;
RD: radial diffusivity;
ROI: region of interest;
SSIM: structural similarity index

## 1. Introduction

Since the first depiction of the diffusion process in the human brain significant conceptual and methodological developments have been applied to diffusion MRI ^1^, leading to the widespread use of various MRI techniques based on this phenomenon, such as diffusion-weighted imaging, diffusion tensor imaging (DTI, ^2,3^), intravascular incoherent motion (IVIM) imaging ^4^, diffusion kurtosis imaging ^5,6^ or diffusion spectrum imaging (DSI, ^7^). DTI offers increased sophistication over diffusion-weighted MRI since it provides information about both the magnitude and orientation of the anisotropic diffusion in tissues, consequently allowing the calculation of the magnitude of diffusion anisotropy, and the parallel and perpendicular diffusivity. According to basic experiments, such parameters in the human brain reflect the underlying axonal membrane microstructure ^8^, correlate with myelination patterns ^9^, can serve as group-level markers of various brain pathologies ^1,10-16^, and allow more complex image post processing approaches, such as fiber tractography ^17,18^. Diffusion MRI approaches are therefore widely regarded as nascent methods for characterising the human brain from its tissue microstructure to its network-level connectional architecture, referred to as the connectome ^19^. The usability of DTI, owing to recent advancements in fast imaging sequence development, can be extended to the earliest point of the human lifespan: before birth, as early as the second trimester of gestation ^20^. Initial *in utero* DTI studies have revealed how commissural, projection and association fibers emerge in the living human fetus ^21,22^, detected pathological, ectopic fibers ^23-25^, and shown surprisingly good agreement with similar, *post mortem* fetal MRI studies ^26^.

As DTI was applied to ever more demanding experimental and clinical scenarios, it faced numerous tests of reproducibility, resulting from the increased sensitivity of DTI and other echo-planar imaging (EPI) sequences towards artifacts. Such experiments are of key importance when we consider the so-called secondary maps – such as the fractional anisotropy – quantitative makers of tissue microstructure in normal and pathological conditions. Although DTI, especially in adults and children, can be used to calculate the scalar metrics of diffusion with high confidence ^27-30^, and tractography has also been shown to be reproducible under constant conditions ^31-35^, it is now clear that differences in the scanner type or imaging site ^36-40^, the parameter settings of the sequence and possibly other factors greatly hinder the comparability across research sites. Fetal DTI is further complicated by excessive motion and more pronounced susceptibility artifacts because of the heterogeneous chemical composition of the maternal organs surrounding the fetal head.

Despite the technical challenges associated with *in utero* DTI, it is currently the only clinically viable imaging modality capable of visualizing the developing white matter during the second and third trimesters of gestation. The diffusion tensor approach provides added value to that of diffusion-weighted imaging in that the fractional anisotropy, the eigenvalues and orientations of the tensor may reflect many, thus-far not thoroughly studied, physiological and microstructural attributes of fetal nerve tissue. DTI and tractography before birth therefore represent important approaches for basic and clinical neuroscience research, especially since our current understanding of the transient morphology of the emerging human brain pathways and its vulnerability during “risk periods” of development are based on postmortem histology and imaging in a limited number of subjects. In vivo MRI validations of advanced early brain imaging data are only available from the 24-26^th^ weeks in extremely preterm neonates, which is a suboptimal model for the characterization of white matter development. In contrast to the post mortem, animal investigations and postnatal data of preterm infants, a prenatal DTI based work-up augmented with tractography can provide new insight into fiber development in normally developing fetuses or reveal the *in utero* trajectory of pathological fiber development. Such a rich parameter set may not only be viewed as a fingerprint of normal and pathological development on the case-level basis, but as potential candidates for prenatal biomarkers, and thus they can hold the clue to predicting the outcome of pregnancy and the neurological outcome after birth.

This endeavor, however, should be preceded by validation experiments aiming to characterise the reliability of the quantitative metrics of diffusion, and to identify the possible factors that can confound their applicability as markers of disease in fetal MRI. Our purpose, therefore, was to demonstrate the within-subject reproducibility of *in utero* DTI in a clinical cohort of fetuses with unaffected and pathological brain development taken from a clinical sample, and to evaluate how scanner field strength, fetal age, fetal motion patterns or the presence of pathology affect the reproducibility of the DTI derived metrics of diffusion.

## 2. Methods

### 2.1 Study population

The study population consisted of fetuses for which fetal MRI was clinically indicated. The general clinical indication for fetal MRI is in conditions where the combined accuracy of MRI and ultrasound is higher than with the ultrasound alone. In our study, the indication to perform fetal MRI was to rule out or confirm brain or lung pathologies or for post-operative follow-up after open fetal correction for spina bifida. Demographic data, clinical indication and the summary of neuroimaging findings are summarized in **Table 1**. As part of the routine clinical protocol, fetal MRI included (1) multi-planar structural MRI examinations with T2-weighted sequences of the whole fetus, fetal brain and the placenta, (2) T1-weighted and echo planar diffusion-weighted sequences to rule out intracranial hemorrhage and/or blood breakdown products, and (3) DTI, for which only the non-processed, trace-weighted *isotropic image* was used in clinical diagnostics.

**Table 1.**
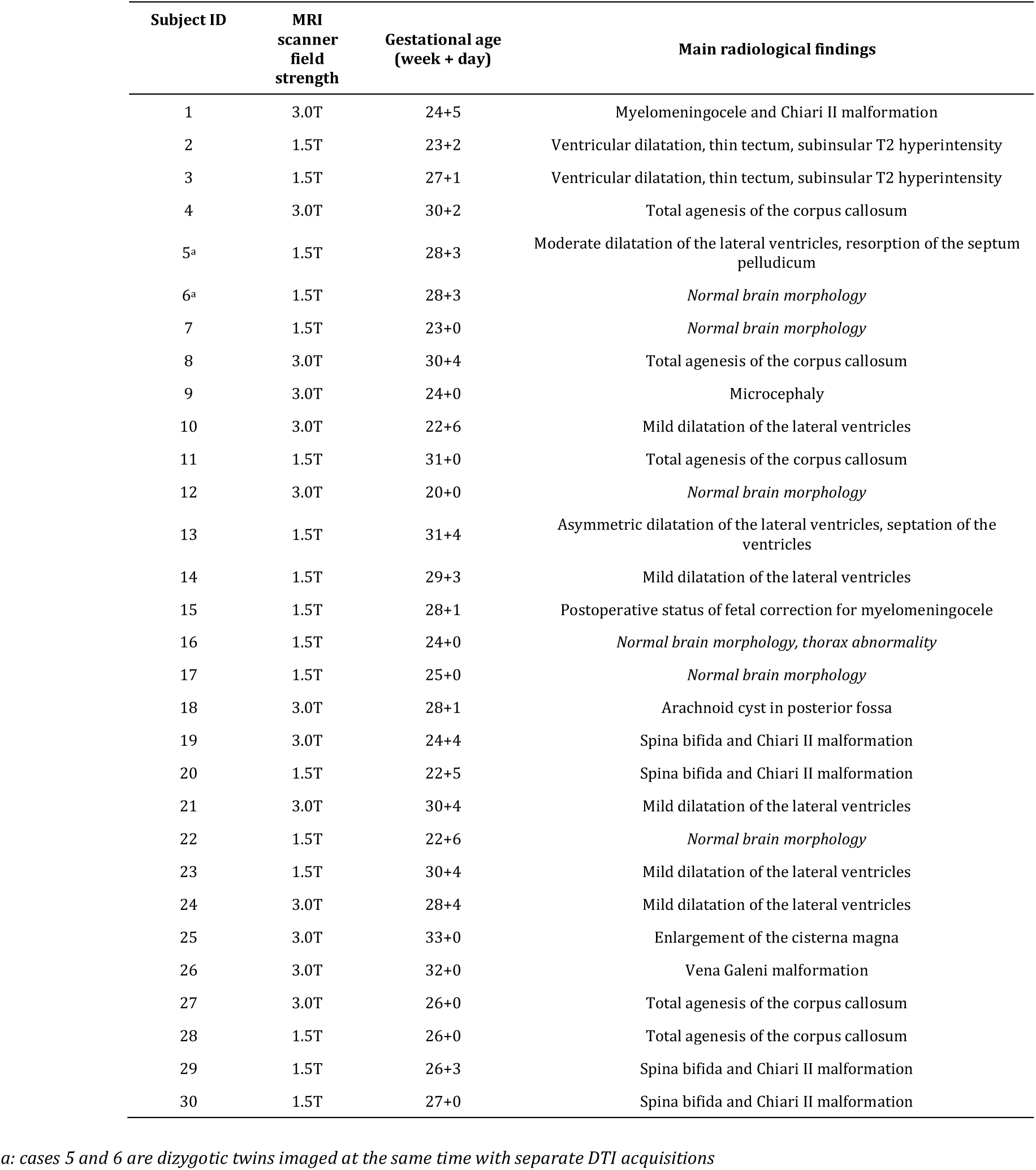
*Demographic data of the study population.*

Inclusion criteria were: fetal age equal to or higher than 20 weeks of gestation based on ultrasound report prior to the MRI, availability of three repeated-session DTI scans with feasible image quality and with only a few motion-corrupted time frames (<5). During the study period between January 2016 and January 2017, we enrolled 30 fetuses in the study, with a mean gestational age of 27 ± 3.3 (range: 20-33) weeks. Of 30 fetuses, 6 had normal brain development according to the ultrasonography and fetal MRI reports. The mothers gave written, informed consent for use of their clinical data for research purposes prior to the examination, and the research was conducted according to the principles expressed in the Declaration of Helsinki. The study was approved by the regional ethical committee.

### 2.2 Fetal diffusion tensor imaging protocol

As a part of clinical routine, fetal MRI was performed on two different clinical MRI systems, one with a field strength of 1.5 T and one of 3.0 T (MR450 and MR750, GE Healthcare, Milwaukee, WI, USA). The assignment of the cases to either of these scanners was not controlled in the current study, and was based on the availability of free scanner time. Pregnant women were examined in the supine position (feet first), and no contrast agents or sedatives were administered. In order to obtain optimal MR signal, the coil was readjusted depending on the position of the fetal head during the imaging procedure. DTI scans followed the structural, T2-weighted images, and three repeated DTI sessions were performed consecutively. Axial slices were positioned orthogonal to the fetal brainstem. The basic settings of the DTI protocol were identical for the two MR scanners used. For DTI acquisitions, an axial, single-shot, echo planar imaging sequence was used with a TR of 2200 ms, a TE of 63 ms, acquisition matrix of 112 * 112 re-sampled to 256 * 256, a voxel size of 1 * 1 mm, and a slice thickness 3 to 5 mm without a gap or interleaved slices. Depending on the size of the fetal brain and gestational age, 10-18 slices were acquired, covering the whole brain from the brainstem to the convexities. Images were acquired using SENSE and a pseudo-receive bandwidth of 33 Hz/pixel along the phase encode dimension. For each of the three repeated DTI scans, 15 non-collinear diffusion-weighted magnetic pulsed gradients were used with a b value of 700 s/mm^2^ and one B_0_ image without diffusion weighting was also collected. Fetal DTI data were anonymized and transferred to image-processing workstations in DICOM format.

### 2.3 Standard and fetal-specific post processing of DTI data

After transferring the MRI data to the image processing workstations the following analysis steps were performed: (1) manual masking of the fetal brain volume to separate non-brain tissue, (2) fetal-specific postprocessing including motion correction of DTI frames, (3) standard diffusion tensor estimation and calculation of the scalar measures of diffusion, (4) detection of white matter structures and ROI placement on major fiber bundles using colorized fractional anisotropy and fractional anisotropy images, (5) qualitative and quantitative analysis of image reproducibility.

Due to the presumed confounding effects of fetal head and maternal respiratory motion on the quality of the reconstructed DTI data, we corrected the images for spurious fetal movements. First, the fetal brain was manually masked on the first image frame of each of DTI scan. To ensure that the brain’s borders remain precisely within the mask despite the possible movements in the consecutive images, a machine learning algorithm was utilized that propagated this mask along the time dimension in each scan. This was followed by the co-registration of each masked image frame to the first reference image of the scan to achieve identical orientation and good anatomical overlap. The fetal-specific image processing algorithm is described in detail in the **Supplementary Document.** After the re-orientation of the raw diffusion-weighted images to reduce the effects of fetal head motion, we performed standard diffusion tensor estimation using the *dtifit* command in the FDT toolkit of the FSL software package. For the diffusion estimation, the b-vector orientations were corrected for the estimated head rotation, and a weighted least squares approach was used. We calculated the fractional anisotropy (FA), mean diffusivity (MD), axial diffusivity (AD) and radial diffusivity (RD) from the tensor datasets using the known general equations from the literature ^10^. For further post-processing steps, the mean image of all the non-B_0_ image frames was calculated for each scan; we will refer to this image as the “mean DWI”.

The next steps of the analysis included the qualitative assessment of fiber visibility on each repeated scans, ROI placement and the voxel-wise comparison of values across repeated scans. For the last two steps, good anatomical overlap between the repeated imaging within each subject was required. Based on the masked images of the mean DWI, two transformation matrices were determined that transformed the second and third repeated scans to the space of the first DTI of each subject. This linear co-registration was performed by the *flirt* command in the FSL software package, which utilized the least square differences as the cost function and optimized the transformation using 6 degrees of freedom. These matrices were used to re-sample the repeated FA, MD and AD scalar maps to enable the comparison of voxel-wise values and ROI-based values. An overview of our study work-flow is illustrated in **Figure 1.**

**Figure 1.**
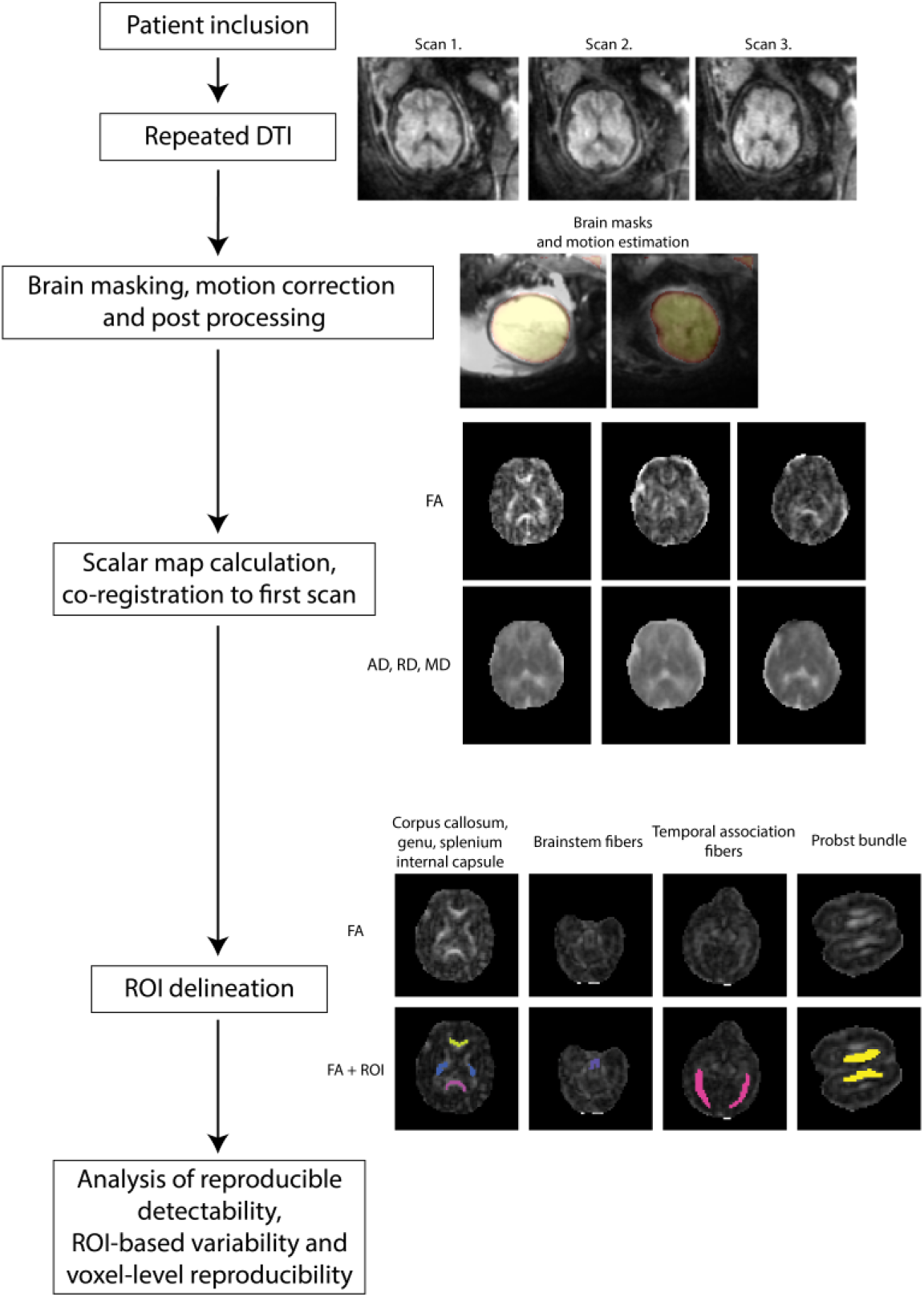
Testing the reproducibility of in utero DTI: overview of the study work-flow.

### 2.4 Definition of white matter structures

Due to the ongoing development of white matter structures during mid-gestation and the technical difficulties of imaging, we selected five prenatally visible major fiber bundles and evaluated whether they can be detected reproducibly across repeated DTI scans. A structure was classified detectable if a physician with experience in fetal anatomy could see it on at least two axial slices of the fractional anisotropy images. We evaluated the (1) anterior and (2) posterior part (genu and splenium) of the corpus callosum at the level of the third ventricle, (3) the bilateral internal capsule, posterior limb, (4) the brainstem fibers consisting of projection fibers including the corticospinal tract and the medial lemnisci at the level of the pons to mesencephalon and the (5) fiber pathways that surround the occipital horn of the lateral ventricles posterolaterally. This latter structure comprised the inferior fronto-occipital fascicle, the optic and acoustic radiations and the subcortical fibers of the temporal and occipital lobe, and therefore we used the term “temporooccipital association fibers” in our study. In 3 cases with corpus callosum agenesis, the ectopic bundle of Probst was evaluated instead of the callosal fibers.

### 2.5 Analysis of white matter structure detectability

In our study, reproducibility was defined using two concepts: the ability to detect major white matter structures in repeated scans within one MRI session in each individual, and the variability of quantitative measurements of a given structure across the repeated scans. We tested the reproducibility of *in utero* DTI using the following qualitative and quantitative approaches: (1) the observer based visibility of white matter structures on FA maps, (2) reproducibility of DTI derived measures in anatomically important regions of interests (ROIs) and the (3) voxel-wise variability, image correlation and image similarity measured over the whole fetal brain. During the qualitative analysis of tract visibility and reproducibility, the following categories were assigned to each tract in each subject. A tract was classified “not visible”, if it cannot be seen on any of the repeated scans, “visible, not reproducible” if it was only seen on one scan, “moderately reproducible” if it was visible on two scans, “highly reproducible” if it was visible on all three scans. For each subject, the occurrence was also calculated, which captured how often a particular tract is visible during the repeated scans (%).

### 2.6 Analysis of repeatability

After this qualitative evaluation, the FA, MD, RD and AD values were measured in each ROI where the tract was detectable on the FA images. Variability was quantified as the coefficient of variation (CV) in percent units across repeated scans using the following equation:

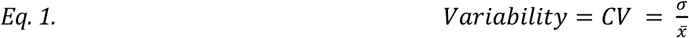

where σ is the standard deviation across the repeated scans, and 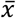 is the mean value across the repeated scans, σ is defined as:

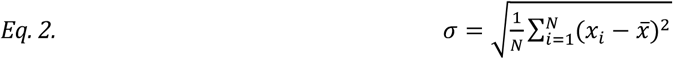

*where N is the number of repetitions in which the tract is visible, 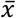 is the mean value across the repeated scans, *x_i_* is the i*^*th*^ *value..*

The voxel-wise reproducibility of images tested whether values across the entire brain could be reproduced. For this analysis step, the images were co-registered to the first scan of each subject in order to achieve anatomical correspondence. A Gaussian filter was applied to the images with a full width at half maximum (FWHM) of 2 mm to correct for small remaining sampling errors due to the imperfect overlap between the scans.

Three parameters were used to characterise the reproducibility of each DTI metric on a per-voxel basis. First, we calculated the standard deviation of DTI metrics using Eq. 2 and then averaged the values over all brain voxels. The image correlation was calculated using the Pearson product-moment correlation coefficient, averaged over all brain voxels. We then calculated the Structural Similarity (SSIM) Index for each image pair ^41^. SSIM is a quality assessment index, which is based on the computation of three terms, namely the luminance term, the contrast term and the structural term. The overall index is a multiplicative combination of the three terms:

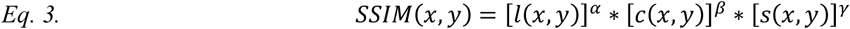

*where for a given pixel x,y, l, c and s are the luminance term, the contrast term and the structural term, and*

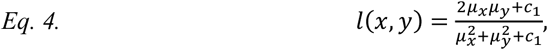

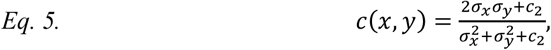

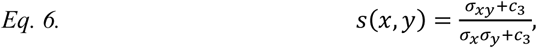

*where μ*_*x*_*, μ*_*y*_*, σ*_*x*_*,σ*_*y*_*, and σ*_*xy*_ *are the local means, standard deviations, and cross-covariance for images x, y.*

We evaluated whether the gestational age of the fetus, the field strength of the scanner, the motion correction, the magnitude of fetal head movement and the position of the fetal head influenced the reproducibility measures. The influence of these factors on the white matter structure detectability was tested by comparing the reproducibility measures between the groups formed by the categorical variables, such as scanner field strength, using the Wilcoxon rank sum test. The influence of continuous variables, such as the gestational age and fetal head movement on reproducibility was tested by linear regression, and outlier effects were addressed using the least squares fitting approach in Matlab for Windows R2014 (Mathworks Inc, Natick, MA, USA).

## 3. Results

### 3.1 Visibility of white matter structures across repeated scans

The corpus callosum was investigated in 25 cases; in the remaining 5 cases, due to the total agenesis of the corpus callosum, ectopic bundles were examined in place of the corpus callosum. The genu and splenium of the corpus callosum were the most commonly visible structures in our study, demonstrating a moderate to high reproducibility in 76% (genu) and 64% (splenium) of the cases, meaning that the structures were detectable in at least 2 of the 3 repeated DTI scans. However, the central part of the corpus callosum was usually not visible, the only exceptions were larger fetuses in whom the thicker corpus callosum was generally easier to delineate. Without motion correction, the visibility of these structures was considerably lower: 60% and 44%, respectively. The bilateral internal capsule’s posterior limbs were successfully visualized in 93.3% of the cases in at least one of the repeated DTI scans, although in 33.3% of the fetuses this structure was not reproducible and was only seen in one of the three scans, while in 40% of the cases it was seen in all of the three repeated scans. Without correcting the images for fetal motion, the internal capsule, posterior limb was highly reproducible only in 20% of the cases, and was not visualized in 9 cases. The temporooccipital association fibers, which include different pathways lateral to the ventricles in the temporal and occipital lobe, were moderately to highly reproducible (moderate reproducibility: 16.7%, high reproducibility: 43.3% of the cases). The least detectable structures in our analysis were the brainstem fibers, which were only seen repeatedly in 40% of the cases and high reproducibility was only reported for 4 fetuses (13.3%). The ectopic Probst bundles were seen repeatedly in 4 of the 5 CCA cases. The qualitative visibility results after fetal-specific image processing for each tract are summarized in **Table 2.**

**Table 2.** *Visibility of major white matter structures in utero across the repeated DTI scans.* *A tract was classified “not visible”, if it cannot be seen on any of the repeated scans, “not reproducible” if it was only seen on one scan, “moderately reproducible” if it was visible on two of the three scans, “highly reproducible” if it was visible on all three scans. Frequency means the occurrence and detectability of the tract in an individual across the repeated scans (0-100%).*

| Name of structure | Not visible, %<br>(number of subjects) | Not reproducible, %<br>(number of subjects) | Moderately reproducible, %<br>(number of subjects) | Highly reproducible, %<br>(number of subjects) | Frequency<br>(%, mean ± SD, min-max) |
| --- | --- | --- | --- | --- | --- |
| Corpus callosum, genu | 16%<br>(4 / 25) | 8%<br>(2 / 25) | 28%<br>(7 / 25) | 48%<br>(12 / 25) | 69.3 ± 37.1% |
| Corpus callosum, splenium | 20%<br>(5 / 25) | 16%<br>(4 / 25) | 28%<br>(7 / 25) | 36%<br>(9 / 25) | 60 ± 38.5% |
| Internal capsule, posterior limb | 6.67%<br>(2 / 30) | 33.3%<br>(10 / 30) | 20%<br>(6 / 30) | 40%<br>(12 / 30) | 64.4 ± 33.8% |
| Brainstem fibers | 30%<br>(9 / 30) | 30%<br>(9 / 30) | 26.7%<br>(8 / 30) | 13.3%<br>(4 / 30) | 41.1 ± 34.7% |
| Temporal and occipital association fibers | 20%<br>(6 / 30) | 20%<br>(6 / 30) | 16.7%<br>(5 / 30) | 43.3%<br>(13 / 30) | 61.1 ± 40.2% |
| Probst’s bundle | 0%<br>(0 / 5) | 20%<br>(1 / 5) | 20%<br>(1 / 5) | 60%<br>(3 / 5) | 66.7 ± 33.3% |

### 3.2 Reproducibility of DTI measures

The results of the reproducibility analysis for each white matter structure are summarized in **Figure 2.** The variability of DTI measures across repeated scans in the individual fetuses spanned a large interval from 0.2% to 40.4% with several outliers, e.g. individual cases demonstrating a high variability for all tracts. The variability of values for a given structure averaged over the population was between 3% (Variability of axial diffusivity of the Probst bundle) and 14.6% (Variability of FA, brainstem). Generally, the mean diffusivity, which is rotationally invariant, was less variable, while the FA was the most variable of the examined DTI metrics. The pathological Probst bundle showed considerably lower variability (3% -7.5%) compared to the other structures. The tracts with the lowest variability after the Probst bundle were the temporal association fibers, the splenium of the corpus callosum and the internal capsule, posterior limb. While the genu and splenium of the corpus callosum were the two most frequently visible tracts across repeated scans, the variability of the DTI derived metrics was relatively high in these compared to the other structures..

**Figure 2.**
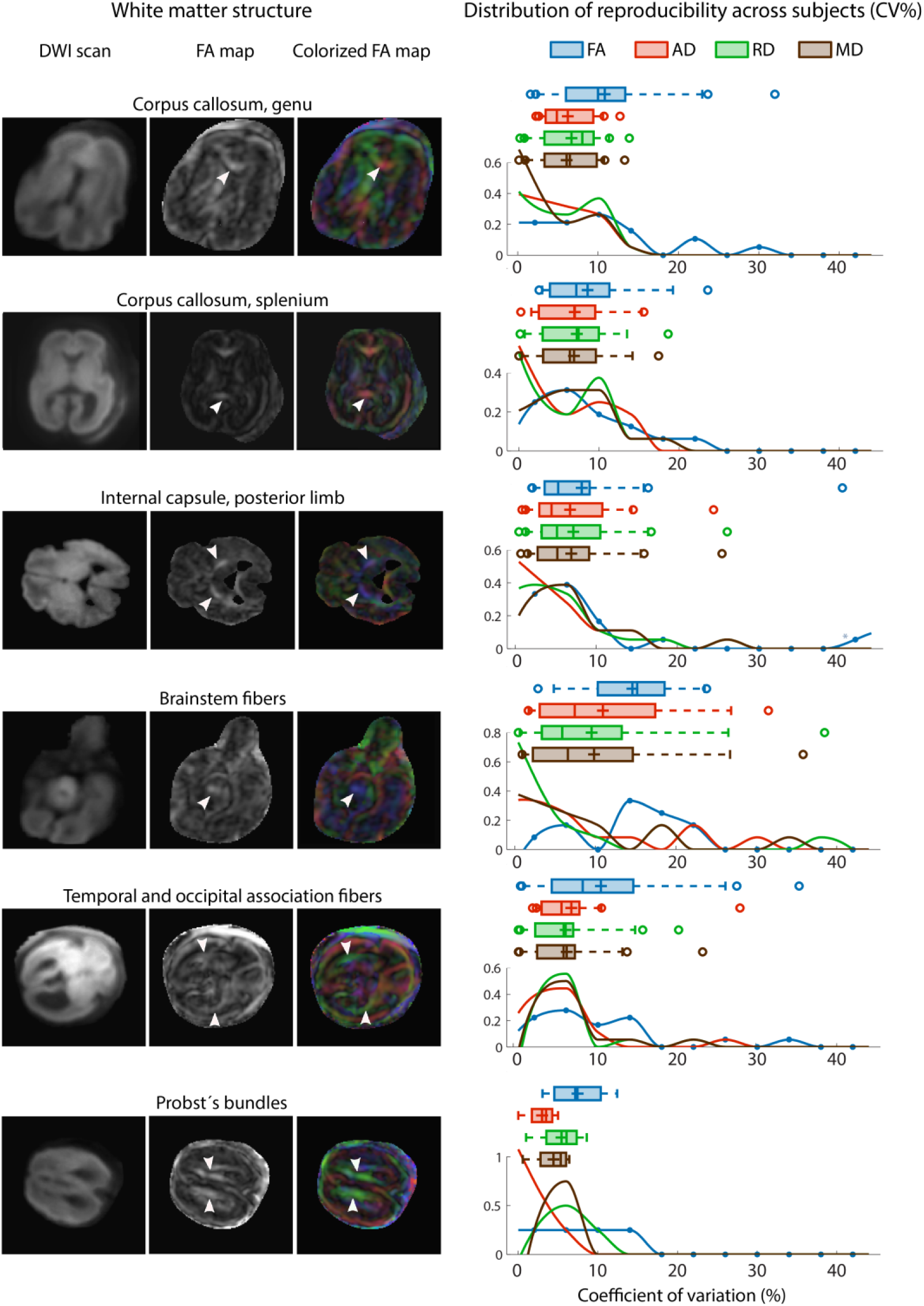
*Reproducibility of DTI derived metrics in selected white matter structures.* *We illustrated each of the evaluated white matter structures on the mean (trace-weighted) diffusion image, on the fractional anisotropy and on the colorized fractional anisotropy maps (left column). The reproducibility of the fractional anisotropy (FA), axial diffusivity (AD), radial diffusivity (RD) and the mean diffusivity (MD) is given as the coefficient of variability (CV%) across the repeated scans. We illustrate the distribution of CV% across the study population for each white matter structure separately (right column). In each diagram, the Whisker plots represent the interquartile range (bar), mean value (horizontal line), median value (cross), outliers (circles), while the histogram demonstrates the distribution of CV% values for each DTI metric separately.*

The results for each white matter structure and each of the four investigated DTI derived metrics are summarized in **Table 3**. We also investigated the distribution of variability across the population, which gives a better representation of the actual variability than the mean and standard deviations, due to the high skew of the distribution. **Figure 2** shows the frequency histograms of the variability of values and demonstrates that a few fetuses had exceedingly high variability across repeated scans, (in some cases and structures more than 20-25%), most likely due to acquisition related errors. With motion correction, the variability of values stayed under 10% for the majority of the subjects. The distribution of variability values also showed less scatter using motion correction and contained fewer extreme values, meaning that the motion correction procedure can ameliorate some of the effects leading to increased variability, at last in some cases.

**Table 3.** *Variability of fetal DTI measurements in different white matter structures across repeated scans.* *Variability is given as coefficient of variation of DTI metrics across the repeated scans, in % units. For each anatomical region, the following parameters are calculated: population mean ± standard deviation, value range and number of subjects (n) with suitable repeated measurement of the given ROI. Variability was calculated for only the subjects in whom the given structure was recognizable in at least two of the three repeated scans.*

| Name of region | Variability (%) of FA values | Variability (%) of MD values | Variability (%) of AD values | Variability (%) of RD values |
| --- | --- | --- | --- | --- |
| Corpus callosum, genu | **10.8 ± 7.7**<br>(1.6 - 31.9)<br>n = 19 | **6.1 ± 3.8**<br>(0.2 - 13.3)<br>n = 19 | **6.2 ± 3.29**<br>(2.3 - 12.7)<br>n = 19 | **6.7 ± 4**<br>(0.29 - 13.9)<br>n = 15 |
| Corpus callosum, splenium | **8.6 ± 6.1**<br>(2.5 - 23.6)<br>n = 16 | **6.9 ± 4.9**<br>(0.2 - 17.4)<br>n = 16 | **7 ± 4.9**<br>(0.2 - 15.6)<br>n = 16 | **7.2 ± 4.9**<br>(0.15 - 18.6)<br>n = 16 |
| Internal capsule, posterior limb | **8.2 ± 8.8**<br>(2.1 - 40.4)<br>n = 18 | **6.9 ± 6.1**<br>(0.7 - 25.5)<br>n = 18 | **6.8 ± 6**<br>(0.8 - 24.5)<br>n = 18 | **7.2 ± 6.5**<br>(0.5 - 26.1)<br>n = 18 |
| Brainstem fibers | **14.6 ± 6.5**<br>(2.9 - 23.8)<br>n = 12 | **9.8 ± 10.4**<br>(0.9 - 35.8)<br>n = 12 | **10.9 ± 9.7**<br>(1.7 - 31.5)<br>n = 12 | **9.5 ± 10.8**<br>(0.4 - 38.5)<br>n = 12 |
| Temporal and occipital association fibers | **10.8 ± 9**<br>(0.9 - 35.3)<br>n = 18 | **6.6 ± 5.3**<br>(0.7 - 23.4)<br>n = 18 | **7.2 ± 5.8**<br>(2.4 - 28)<br>n = 18 | **6.5 ± 4.9**<br>(0.6 - 20.5)<br>n = 18 |
| Probst’s bundle | **7.5 ± 3.9**<br>(3.1 - 12.5)<br>n = 4 | **4.4 ± 2.62**<br>(0.6 - 6.7)<br>n = 4 | **3 ± 2.1**<br>(2.4 - 28)<br>n = 4 | **5.5 ± 3.2**<br>(1 - 8.6)<br>n = 4 |

### 3.3 *Image variability, correlation and similarity*

In **Figure 3** we summarized the results of the voxel-wise analysis of value variability, voxel-to-voxel image correlations and structural similarity averaged for the entire brain. During the ROI based analysis, we concluded that the distribution of reproducibility is highly skewed due to outliers, i.e. fetuses whose DTI data are profoundly corrupted by artefacts in whom variability is very high. This observation was more pronounced for the whole-brain voxel-wise analysis, most likely due to the fact that the DTI metrics calculated within the gray matter and the ventricular system show inherently higher variability. Therefore we decided to exclude 4 fetuses from this analysis, in which the mean framewise displacement was the highest (threshold, 90^th^ percentile: 12 mm).

**Figure 3.**
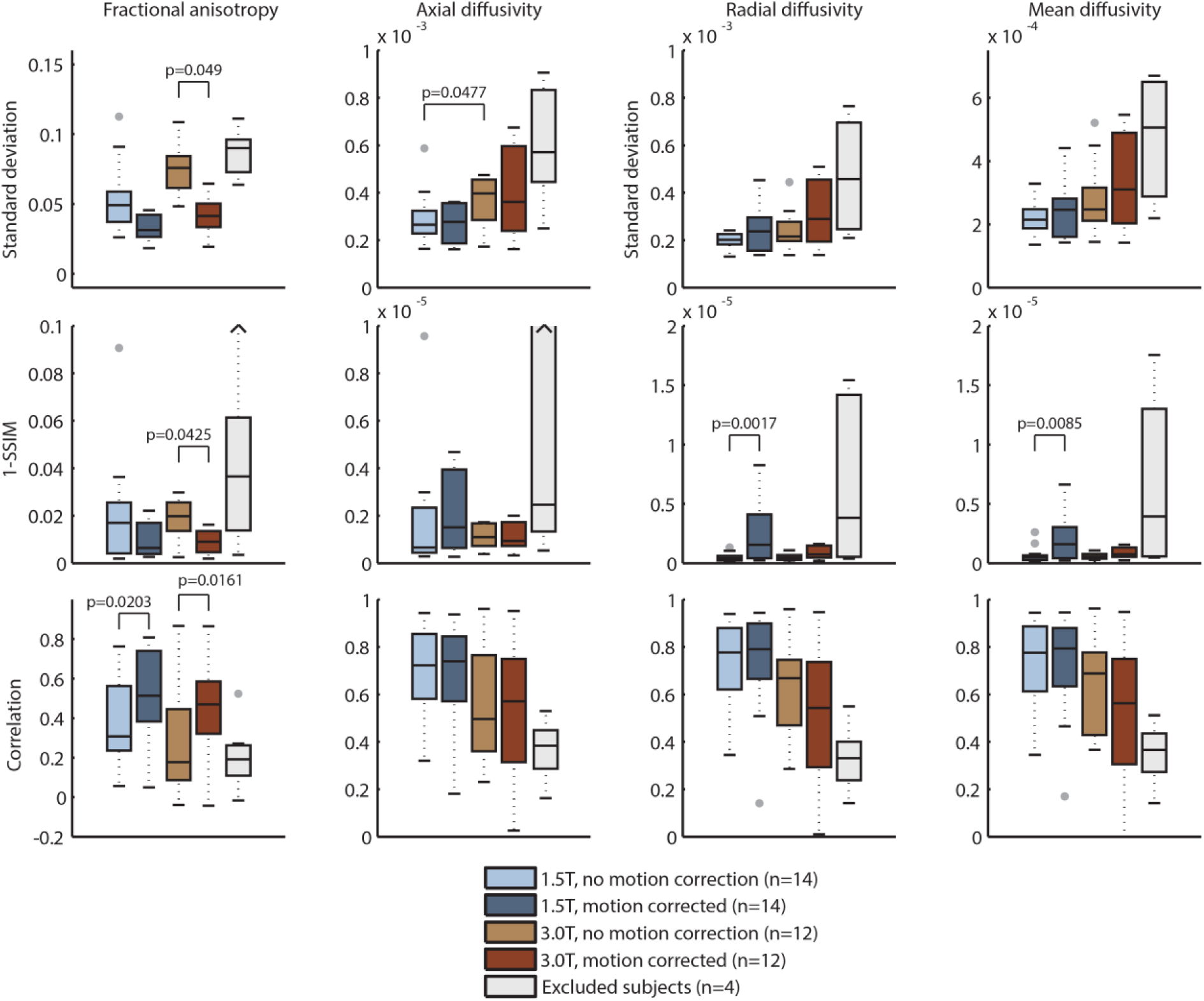
*Voxel-wise reproducibility of DTI derived values based on whole-brain analysis.* *We calculated the standard deviation, the image correlation and the structural similarity index (depicted as 1-SSIM) across scans. Each Whisker-plot demonstrates the distribution of these reproducibility metrics in cases that underwent fetal DTI on 1.5T or 3.0T, with and without fetal specific image processing including motion correction. The fifth plot demonstrates the variability of values in subjects that were excluded based on the criterion of excessive head motion during the DTI scan.*

After motion correction and excluding the cases with the highest motion-related artifacts, the average correlation between DTI derived metrics across the repeated images was high, FA: R=0.494 ± 0.23 (–0.0432 – 0.862), AD: R=0.598 ± 0.258 (0.02 – 0.951), RD: R=0.638 ± 0.269 (0.01 – 0.947) and MD: R=0.632 ± 0.274 (0.076 – 0.949). Structural similarity was also high for all cases after motion correction, ranging from 0.909 to 0.998.

The motion correction procedure had a significant effect and resulted in lower standard deviation of FA values across the 3.0T scans (p=0.0049), higher image correlation of the FA maps (1.5T cases: p=0.0203, 3.0T cases: p=0.0161) and higher structural similarity of the FA images (p=0.0425), tested using paired Wilcoxon signed rank tests. Controversially, the motion corrected RD and MD images had lower structural similarity index on 1.5T (p=0.0017 and p=0.0085, respectively).

Scanner field strength only had a significant effect on the non-motion corrected axial diffusivity images on 1.5T. The standard deviation of the AD maps was significantly higher on 3.0T meaning lower reproducibility, p=0.0477.

### 3.4 Factors influencing the reproducibility

Our study population included fetuses with normally appearing brain development and pathological cases. The presence of pathology affected the visibility of all fiber structures, they were less detectable in the pathological group, as indicated by the lower overall visibility score (normal brain development: 11.29 ± 5.56, pathologies: 7.83 ± 3.94). Fetuses with normally developing brains had lower variability of the FA of the temporoocipital association fibers and the MD of the splenium of the corpus callosum.

The effects of gestational age were pronounced the images that were not corrected for motion. Strong, positive correlations were evident between the gestational age of the fetus and the variability of FA, MD, RD and AD values of the brainstem and internal capsule. After fetal specific motion correction, the correlations between gestational age and variability were only moderate (R^2^<0.25). Interestingly, larger fetuses tend to have larger reproducibility after the robust linear regression analysis, and the split between small (second trimester) and larger (third trimester) fetuses revealed that second trimester fetuses may have higher variability overall for the DTI derived measures.

Two MRI devices from the same vendor were used, which makes it possible to investigate the differences in reproducibility of DTI metrics on 1.5 T and 3.0 T field strengths, with nearly identical image acquisition parameters. The variability of the FA, MD and RD values of the temporooccipital association fibers were approximately twice as high on 3.0 as on 1.5T, meaning lower reproducibility for these structures. (**Table 4**).

**Table 4.** *Factors significantly influencing the visibility of structures and the variability of in utero DTI measures across repeated scans.* *We evaluated whether gestation age, fetal head movement, presence of pathology or scanner field strength influences the visibility of tracts or the various variability measurements. R: Pearson’s product moment correlation coefficient, p: statistical significance of a Wilcoxon rank sum test. FA: fractional anisotropy, GW: gestational weeks, MD: mean diffusivity, SD: standard deviation, SSE: sum of squares of error.*

| Reproducibility parameter | Factor influencing reproducibility | Statistical values (Adjusted-R<sup>2</sup>, or p value) | Values across groups (mean ± SD, range) |  |
| --- | --- | --- | --- | --- |
| Variability of FA values, temporooccipital association fibers | Scanner field strength | p=0.020 | 1.5 T<br>6.9% ± 5.0% ( 0.9% - 14.9%) | 3.0 T<br>17.1% ± 10.6% ( 5.4% - 35.3%) |
| Variability of AD values, temporooccipital association fibers | Scanner field strength | p=0.011 | 1.5 T<br>4.8% ± 1.9% ( 2.4% - 8.2%) | 3.0 T<br>10.9% ± 7.9% ( 4.5% - 28.0%) |
| Variability of RD values, temporooccipital association fibers | Scanner field strength | p=0.011 | 1.5 T<br>4.2% ± 2.7% ( 0.6% - 7.4%) | 3.0 T<br>10.1% ± 5.8% ( 5.5% - 20.4%) |
| Overall visibility score of white matter structures | Presence of pathology | p=0.039 | Pathology present<br>7.83 ± 3.94 ( 1.00 - 13.00) | Unaffected brain development<br>11.29 ± 5.56 ( 1.00 - 15.00) |
| Variability of FA values, temporooccipital association fibers | Presence of pathology | p=0.035 | Pathology present<br>13.2% ± 9.5% ( 1.0% - 35.3%) | Unaffected brain development<br>4.8% ± 2.9% ( 0.9% - 8.1%) |
| Variability of MD values, corpus callosum, splenium | Presence of pathology | p=0.042 | Pathology present<br>8.9% ± 4.9% ( 0.0% - 17.4%) | Unaffected brain development<br>3.6% ± 2.6% ( 0.4% - 7.1%) |
| Visibility of corpus callosum, splenium | Presence of pathology | p=0.023 | Pathology present<br>0.41 ± 0.39 ( 0.00 - 1.00) | Unaffected brain development<br>0.81 ± 0.38 ( 0.00 - 1.00) |
| Variability of MD values, internal capsule, posterior limb | Head movement of fetus | R<sup>2</sup>=0.612 |  |  |
| Variability of AD values, internal capsule, posterior limb | Head movement of fetus | R<sup>2</sup>=0.605 |  |  |
| Variability of RD values, internal capsule, posterior limb | Head movement of fetus | R<sup>2</sup>=0.531 |  |  |
| Visibility of corpus callosum, genu | Head movement of fetus | R<sup>2</sup>=0.255 |  |  |

The factor which seemed to influence the reproducibility measures the most was the mean displacement of the fetal head during the repeated scans. This parameter can be measured during the motion correction step and was expressed in mm scale (mean framewise displacement). Generally, moderate to strong correlations were found between fetal head motion and nearly all of the variability measures. In **Table 4** we only report results if the adjusted R-squared value was higher than 0.25 (R≥0.5), which is considered strong correlation according to the rule of thumb by Cohen ^42^, while the results of the entire analysis are detailed in Supplementary Table 3 and 4. The greatest influence of motion on the variability was found before motion correction, and the DTI metrics the FA values were most affected by motion. Specifically, head motion was highly correlated with the variability of the FA values within the internal capsule, posterior limb (correlation with FA variability: R^2^=0.68). After motion correction, four repeatability values were still influenced by the original head motion: the MD, AD and RD values of the internal capsule, posterior limb (R^2^=0.612, 0.605 and 0.531, respectively).

## 4. Discussion

We found that the visibility of major commissural, projection and association fibers in *in utero* DTI maps was moderate to high in 40-76% of the cases, but in a quarter of the cases at least one of the investigated structures was not visible in any of the three repeated scans, which is an important factor to consider for future studies investigating these structures. Unexpectedly, we did not find a close link between fetal age and the visibility of white matter bundles, which would be expected based on the known developmental trajectories of these structures. By the fetal age of 20 postmenstrual weeks, all the reported commissural and projection fibers are already formed ^9^, however, none of the pathways have not yet reached full maturity, and most axons form synapses only at the level of the subplate ^43^. Due to the restricted spatial resolution of the currently available *in utero* DTI, this level of detail of the axonal organization may remain undetectable. Furthermore, the presence of the radial migration pathways and the radial glia renders the subcortical zone highly anisotropic ^44^, contributing to the limited visibility of some white matter structures in fetuses. Since long range association fibers have to “cross” the radial glia perpendicularly within the subcortical white matter, during mid-gestation this network of crossing fibers may restrict the ability of standard clinical DTI protocols to resolve the proper orientation of the principal diffusion vector, leading to a consistent lack of detectability in fetuses at a younger age. However, in the present study, the detectability of major white matter tracts was not significantly related to gestational age, so we assume that the differences in the detectability of major white matter structures arise as a result of technical limitations of the imaging method and do not necessarily reflect the underlying white matter developmental timing.

The Probst bundle was detected in all of the 5 CCA cases, and was reproducibly visible in 4 of these. This structure is identified as a large white matter structure running longitudinally in the horizontal (axial) plane, and which is easily distinguishable from the internal capsule, posterior limb at the level of the centrum semiovale based on the colored fractional anisotropy map (**Figure 2**). The high visibility and high reproducibility of scalar values of this structure, even with such low case numbers, suggest that the Probst bundle may be a promising prenatal maker of callosal agenesis, and in utero DTI may enable the characterisation of how this ectopic bundle is formed ^23-25^. Our findings show that the genu and splenium of the corpus callosum can also be detected with high accuracy on fetal DTI, however, the central part is usually not seen due to its thin cross-section. A viable strategy to increase the visibility of the central parts would be to decrease the slice thickness to 2 mm, which would enable the acquisition of isotropic pixels during DTI, at the cost of roughly doubling the imaging time. In order to confirm the presence of the central part of the corpus callosum, sagittal DTI acquisitions may help, optionally supported by super-resolution sampling techniques ^45^. By the same token, the visibility of the internal capsule, posterior limb may be improved by using coronal acquisitions. The least visible structures in fetal DTI were the brainstem fibers, which were undetectable or not reproducible in 60% of the cases. This is most likely due to the confounding effect of the surrounding CSF spaces, the pulsation of tissues and the general fact that the cross-section of these fibers are small in the axial plane compared to the other investigated white matter structures. The lack of visibility of these structures underscores the need for care when judging the presence of brainstem or pons developmental abnormalities using DTI, as higher spatial and angular resolution is needed to more efficiently image these projection fibers.

In addition to the tract visibility, we also calculated the intrasubject, inter-session variability of the DTI metrics as an important indicator of the clinical usability of fetal DTI. According to our measurements, this aspect of reproducibility depends on the actual anatomical structure: while the corpus callosum and internal capsule had relatively low variability across scans, the brainstem fibers and the temporooccipital association bundles showed generally higher variability. This most likely stems from the fact that structures with less mature fiber structure will have lower anisotropy and therefore lower signal to noise during DTI scans. Furthermore, their size, their proximity to the ventricle system as a confounding factor may also influence the variability of the DTI metrics and the susceptibility of such structures to imaging artifacts.

Before the DTI metrics can be considered as viable prenatal markers of white matter integrity and development, it is important to evaluate their reproducibility based on previously reported data. The reproducibility of DTI derived metrics has been estimated previously in studies of adult volunteers, but those published data cannot be directly compared to our results due to the different study population and the fact that typical fetal DTI protocols differ in design to the optimal adult protocols, due to the requirement for short scan times. Such protocol optimization for fetal DTI usually involves reducing the number of diffusion-weighted gradient directions and the number of image slices. Although six diffusion-weighted magnetic field gradient directions may be sufficient for estimating the FA and MD values ^46^, there is a clear advantage associated with using 15 or 30 directions ^47^. DTI in adults with at least 30 diffusion-weighting directions according to the Jones-30 scheme was reported to be optimal for reproducibly estimating the fractional anisotropy and mean diffusivity ^48^, however, similar investigations for fetuses have not yet been done. The reproducibility of DTI measurements is inferior to that reported in adult volunteers; however, the coefficient of variation (CV) for more than half of the subjects in our study lies in a similar range to that reported in the following adult studies.

Using the 30 direction acquisition scheme, the CVs of the voxel-based and ROI-based diffusion metrics were below ten percent for a similar set of white matter structures to those investigated in our study ^30^, while gray matter structures, i.e. areas of low fractional anisotropy, showed considerably higher variability. Within scanner variability of the FA and MD values of the corpus callosum were reported to be as low as 1.9% and 2.6% ^27^, while Cercignani and colleagues found that histogram-derived metrics of mean diffusivity were highly reproducible (coefficients of variation ranging from 1.72% to 5.56%), as were fractional anisotropy histogram-derived metrics (coefficients of variation ranging from 5.45% to 7.34%) ^36^. A multicenter study reported CVs of 2.2% for ADC, 3.5% for axial diffusivity and 8.7% for FA in the global white matter over time on the same scanner ^39^, while in children, Bonekamp et al. reported a between-scan reproducibility ranging from 2.6% to 4.6% for the FA and from 0.8% to 3.4% for the ADC ^29^. Both observations are consistent with our finding of inferior reproducibility of the direction-dependent metrics, such as FA or AD compared to rotationally invariant measures like the MD. Using 12 diffusion weighting directions, Heiervang and colleagues have shown that inter-session and inter-subject CVs for the FA in the cingulum bundle, pyramidal tracts, optic radiations and genu of the corpus callosum were 5% and 10%, respectively, while inter-session and inter-subject CVs for the MD were below 3% and 8%, respectively ^28^. In a report from Wang et al., out of 60 tractography measurements, 43 showed intersession CVs ≤ 10%, and the most reliable regions were the corpus callosum, cingulum, cerebral peduncles, the uncinate and the arcuate fascicle ^47^.

The effect of differences in scanner design and MRI vendor on the variability of fetal DTI is also an important factor to consider in addition to the intravendor (intrasubject) variability, as the variability in DTI metrics depends considerably on the coil systems, imagers, vendors, and field strengths used for MRI ^38^. In a study by Pfefferbaum et al, DTI measures were reported to show a systematic bias across scanners (CV = 4.5% for FA and CV = 7.5% for trace) ^27^. A study across 5 scanners also reported higher inter-scanner variability than within-scanner variability across time ^39^, and in another similar study, the intra-site CV for FA ranged from 0.8% to 3.0%, while the inter-site CV ranged from 1.0% to 4.1% ^49^. Validation experiments suggest that DTI protocols with sufficiently low coefficient of variation not only allow for the estimation of per-voxel estimates of diffusion, but also the reconstruction of fiber pathways reproducibly using tractography techniques, usually within one study site ^28,31-34,47^. However, we note that the higher interscanner variability (relative to the intrasubject variability) becomes more pronounced during diffusion tractography and causes less comparable results across sites ^40^, and this interscanner variability should also be considered when performing tractography using fetal DTI data.

In the present study, we observed low standard deviations of DTI metrics across the scans, high image correlation and high structural similarity with voxel based whole-brain analysis of the images, and a similar trend was apparent in the ROI-based reproducibility and visibility analysis. While motion correction and the exclusion of severely corrupted data sets improves data reproducibility (**Figure 3**), the 3.0 T DTI acquisitions appear to be inferior in comparison to the 1.5 T data. We observed better reproducibility results when using a voxel-level comparison of the repeated images. The lower variability compared to the ROI approach, high image similarity and correlation for all image voxels, including gray matter and cerebrospinal fluid pixels, may not necessarily reflect improved data usability. Artifacts that have an influence for all brain voxels or large parts of the brain equally can result in inflated per-pixel correlations between scans, while structural similarity may not accurately reflect the absolute variability of values across repeated scans. From previous studies reported in the literature, an intersession DTI metric variability of less than 10% appears to be a ubiquitous finding in adults or in children, which is only comparable to the *in utero* measurements if all fetal scans demonstrating severe artefacts are consistently excluded, leading to a data loss affecting at least 15% to 25% of the subjects.

Our study allows us to estimate the effect of various factors on the reproducibility of *in utero* DTI, including scanner field strength, the presence of pathology and fetal head movements. Significant differences in white matter structure visibility between 1.5 and 3.0 Tesla were found even when fetal-specific image processing was applied. In this case, the variability of diffusion values of the temporooccipital association tracts was higher on the 3.0T data. As 3.0T is being increasingly used for fetal imaging ^50^ due to the higher spatial resolution and better tissue contrast in T2-weighted imaging associated with the higher field strength, this finding raises concerns. A possible explanation for the lower tract reproducibility on 3.0T fetal DTI is that despite the increased SNR and reports of higher reproducibility with higher field strengths in adults ^51^, susceptibility related artifacts arising from the heterogeneous environment surrounding the fetus lead to more pronounced field inhomogeneities, as described in previous reports ^52,53^. To reduce the inter-slice effects arising in EPI acquisitions and the influence of field inhomogeneity, various approaches have been suggested ^54-56^, of which we adapted a normalization of the Z-profile of the intensities on a slice-wise basis, in order to improve our estimate of the diffusion anisotropy. Our study sample is broadly representative of the wider clinical fetal MRI population, in which 0-25% of the cases are typically evaluated as showing normal brain development. This allowed us to judge if DTI metrics from normally developing fetuses have different reproducibility from those estimated in fetuses with intracerebral pathology. The presence of pathology was associated with a worse visibility of fibers overall and the visibility of the splenium of the corpus callosum, which is most likely due to the fact that pathological fetuses often had hydrocephalus, which causes thinning (due to physical compression) of the periventricular fiber structures, resulting in lower visibility when imaged with a large voxel size and slice thickness.

Not surprisingly, the degree of fetal head movement during the scan, the mean framewise displacement, was positively correlated with the variability of nearly all investigated structures. Real motion – i.e., the displacement of the head relative to its environment – can stem from fetal head and trunk movements and maternal breathing, while apparent motion of the brain can also arise from susceptibility related distortions, which are more pronounced *in utero*. These distortions cause spin history artifacts that influence the signal intensity ^56^. Pulsation from the amniotic fluid and surrounding organs, such as the maternal aorta, may further lead to spin dephasing during the diffusion-weighting sequence. The displacement of voxels between the individual diffusion-weighted image frames corrupts the reconstruction of the diffusion tensors and results in lower FA values and incorrect orientations of the calculated eigenvectors ^57^, consistent with our observation that the mean framewise displacement mostly affected the directionally dependent axial diffusivity and the FA.

Our study further enables us to outline two different strategies to tackle the problem of lower reproducibility of current, clinically viable *in utero* DTI sequences. First, the retrospective correction of fetal head movement can be applied to achieve more reproducible reconstruction of the diffusion tensor, for which we used a custom image processing work-flow in our study. Unintentional head motion is a common problem in imaging studies, and various reconstruction approaches have been suggested for fetal functional MRI ^55^, as well as for fetal and neonatal DTI ^58^. A common way to improve image quality is to account for fetal head motion by re-aligning each time frame to a selected reference point ^59^. This realignment step is implemented for common functional MRI processing tools, however, its use in fetal diffusion imaging is not yet established. Retrospective correction of motion was beneficial for our study, which is consistent with findings from the literature ^60^. A particularly promising approach is to reconstruct fetal diffusion-weighted MRI data on regular grids from scattered data ^61^. However, it also appears that fetal head motion reduces reproducibility slightly even if the image frames and diffusion vectors are corrected for motion, and hence further mitigation strategies are needed. Another solution to improve data usability is to collect the DTI in multiple acquisitions spread across the MRI examination. According to our experience, excessive fetal movements that lead to considerable signal loss during DTI may periodically increase or decrease in activity multiple times during a typical, hour-long fetal MRI. It is not possible to forecast such periods, but we can increase the chance for a motion-free DTI by leaving time between the scans. To maximize the clinical output during the “waiting periods”, other clinical sequences designed for observing fetal movements ^62^, or standard, ultra-fast T2-weighted anatomical scans may be acquired. As the fetus may rotate its head significantly in-between the waiting periods, re-orientation of the b-matrix is required during the fetal-specific image post processing (**Supplementary Document**). Repeated DTI has further benefits. Multiple, interleaved B_0_ images allow for the estimation of the noise floor for each pixel, enabling a more robust estimation of the diffusion tensor, such as that using the RESTORE algorithm ^63^. The rotated b-matrix means that images are acquired with slightly different diffusion-weighting orientations, which allows for a subsampling of non-collinear directions to achieve higher angular resolution. Therefore, by utilizing within-subject repeated DTI acquisitions with clinically feasible imaging parameters within a single scanning session, the fiber architecture can be reconstructed in finer, three-dimensional detail, following the “super-resolution” approach ^45,64,65^.

Our prenatal neuroimaging study suffers from a number of important limitations. The statistical power is limited by the exploratory nature of the study, and specifically the small participant group sizes. Increasing the case numbers based on clinical data, especially for fetuses with normal brain development, is challenging due to the time constraints of fetal MRI and the relatively low number of such scans. In addition, the participant group is heterogeneous in terms of gestational age, and the brain anatomy is often affected by various pathologies, such as hydrocephalus, asymmetric lateral ventricle size and associated developmental abnormalities. A further limiting factor originates from the fact that fetal MRI protocols often have to make a compromise between resolution, slice thickness, fast imaging time, patient comfort, reduced SAR, and artefacts arising from fetal motion. Our study is based on a relatively thick-slice DTI protocol, with 15 diffusion weighting directions and a b-factor of 700 s/mm^2^. In particular, the large slice thickness in fetal studies is likely to present a confounding factor when testing reproducibility, since smaller white matter bundles, especially if they run parallel to the imaging plane, may be only partially imaged and good anatomical correspondence between participants is hard to achieve. This effect is more pronounced if the fetus moved or rotated its head perpendicular to the axial plane in any of the image frames, but may be partially mitigated by oversampling the DTI from different directions, as proposed in super-resolution studies for structural and diffusion imaging ^45,61^.

A considerable physiological limitation of the current study is that the exact microstructural correlates of the described white matter bundles are unknown. It is known that in DTI, the majority of the anisotropy stems from the microscopic structure of the axonal membranes ^8^, and to a lesser extent, from the myelin sheath around the axons. Elements of the extracellular matrix, axonal tubules or other, even non-neural structures may only have a limited role in causing anisotropic diffusion. During the fetal age range investigated, however, neither neuronal migration, axonal growth, pathfinding, nor myelination are complete. We cannot rule out the possibility that transient cellular components, such as radial glia may modify the DTI measurements in utero. The clarification of such confounds would require future work with histological work-up in fetal specimens, including further works comparing DTI results *in utero* and *post mortem* and analyzing fiber structures in histological samples^24^.

We conclude that the reproducibility of *in utero* DTI is comparable to that of adult studies only if we exclude the subjects whose images were severely compromised by artefacts. This observation provides a note of caution for studies attempting to use *in utero* DTI as a marker of disease in individual clinical cases: even with scans repeated three times, fiber bundles can remain undetectable and the entire session may be corrupted from artefacts that are not mitigated with image post processing. However, in group studies, *in utero* DTI offer a promising approach for depicting white matter anatomy and measuring microstructural properties of tissue diffusion, and could even allow for the reconstruction of the emerging brain connectivity prenatally. Future directions of fetal imaging research should emphasize the more robust reconstruction of the diffusion tensor based on repeated data and higher angular resolution acquisition schemes. In this endeavor, fetal-specific image processing and repeated scanning is recommended to ensure the detectability of white matter structures.

